# Fentanyl–Xylazine Co-Administration Leads to Sustained Depression of Breathing and Body Temperature Likely Driven by μ-Opioid and α_2_-Adrenergic Pathway Interactions

**DOI:** 10.64898/2026.04.21.719036

**Authors:** Nicole Lynch, Janayna D Lima, Sathyajit Bandaru, Natalia Machado, Satvinder Kaur

## Abstract

**Background and Purpose:** It’s been reported that illicit drug supplies increasingly contain the α_2_-adrenergic agonist, xylazine, alongside fentanyl, yet the pharmacological basis for the greater lethality of this combination remains unclear. Prior research has shown that μ-opioid (Oprm1) receptors, on which fentanyl acts, and α_2_-adrenergic (Adra2a) receptors, on which xylazine acts, are both expressed within brainstem circuits that govern autonomic control, especially the parabrachial (PB) and Kölliker-Fuse (KF) nuclei that regulate respiration. Thus, we propose that co-activation of these inhibitory receptors and their respective pathways could potentiate or additively suppress respiratory and thermoregulatory function.

**Experimental Approach:** Freely behaving C57BL/6J mice received intraperitoneal injections of either saline, fentanyl, xylazine, or fentanyl–xylazine (F+X) solutions. Continuous recordings of respiration using whole-body plethysmography, sleep/wake state using EEG/EMG and body temperature using both infrared thermography, and telemetry were collected for several hours following injection. RNAscope was used to identify Oprm1 and Adra2a expression within PB and KF nuclei.

**Results:** Fentanyl alone produced dose-dependent respiratory depression that was not associated with body temperature changes, whereas the dose we used of xylazine alone had no effect on either respiration or body temperature. In contrast, F+X induced a markedly prolonged (>5 h) reduction in respiratory rate and profound hypothermia lasting 7–8 h, exceeding the effects of either drug alone. Mortality increased to 58.8% following F+X exposure. RNAscope revealed that both Oprm1 and Adra2a receptors are expressed in PB/KF FoxP2-positive neurons, identifying a plausible substrate for convergent inhibitory signaling.

**Implications:** This manuscript provides the first direct experimental evidence that fentanyl and xylazine may interact through convergent μ-opioid and α_2_-adrenergic receptor signaling to produce additive and sustained suppression of respiratory and thermoregulatory function. These findings address a critical mechanistic gap in understanding the disproportionate lethality of fentanyl–xylazine mixtures, an emerging public-health crisis. The work further identifies the PB/KF FoxP2 population as a plausible site of dual-receptor convergence and highlights a previously unrecognized pharmacodynamic interaction with immediate implications for overdose reversal strategies. Given the novelty, mechanistic insight, and translational urgency of these results, rapid dissemination will help accelerate scientific and clinical responses to this evolving threat.

**Graphical Abstract:** Possible convergent μ-opioid (Oprm1) and α_2_-adrenergic (Adra2a) signaling within parabrachial FoxP2-expressing neurons likely produces additive suppression of respiratory and thermoregulatory drive during fentanyl–xylazine co-exposure.
Fentanyl and xylazine engage parallel inhibitory GPCR pathways in Parabrachial/ Kölliker Fuse nucleus (PB/KF) neurons that project to the pre-Bötzinger complex (preBötC) to depress respiratory rhythm and to the dorsomedial hypothalamus (DMH) to blunt thermogenic output. Co-activation of these pathways results in sustained bradypnea, profound hypothermia, and reduced survival, providing a possible mechanistic basis for the increased lethality of fentanyl–xylazine mixtures.

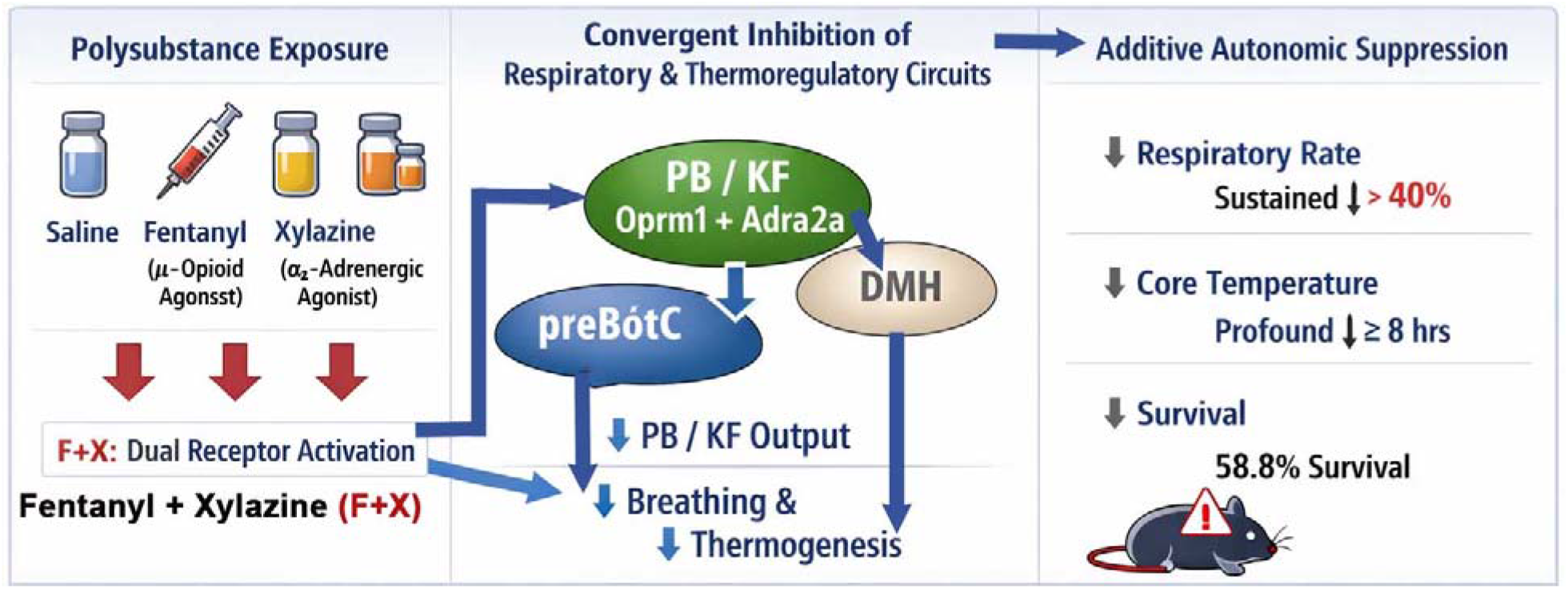

## Introduction

The rapid rise of fentanyl-related overdose deaths in North America has been exacerbated by the emergence of xylazine (X), a veterinary α_2_-adrenergic agonist^1–4^ increasingly detected as an adulterant in illicit opioid supplies^3,5,6^. Xylazine’s infiltration into fentanyl markets has coincided with a marked escalation in overdose fatalities, which is particularly concerning as xylazine produces profound central nervous system depression that is largely insensitive to naloxone^7,8^. Clinical and epidemiological reports consistently show that individuals exposed to fentanyl–xylazine (F+X) mixtures exhibit deeper respiratory depression, prolonged unresponsiveness, and substantially higher fatality rates^3,9^, than those using fentanyl alone^4,10^. Despite these observations, the biological basis for the disproportionate lethality of F+X remains poorly defined.

Fentanyl and xylazine act through distinct inhibitory G-protein–coupled receptors— -opioid receptors (Oprm1) and α2-adrenoceptors (Adra2a), respectively—that regulate overlapping autonomic functions ^1,11–15^. Fentanyl suppresses respiratory rhythm generation and chemosensory drive^16,17^, whereas xylazine at 10mg/kg induces sedation, bradycardia, hypotension, and hypothermia^18,19^. Although these receptor systems are typically studied independently, emerging anatomical evidence indicates that Oprm1 and Adra2a may co-express^20,21^ within key brainstem nuclei governing respiratory rhythmogenesis, arousal, and thermoregulation. Such overlapping localization^22^ suggests that simultaneous activation of these inhibitory pathways could produce convergent or additive suppression of neural circuits essential for maintaining ventilation^23,24^ and core body temperature^25,26^.

Two brainstem hubs are especially poised for such interactions. The pre-Bötzinger complex (preBötC) is required for inspiratory rhythm generation^27^, while the parabrachial (PB) and Kölliker-Fuse (KF) nuclei provide modulatory and arousal-related input that shapes respiratory drive^24,28,29^. MOR (Oprm1)-expressing PB/KF neurons are principal mediators of opioid-induced respiratory depression^30–32^, and PB glutamatergic populations—especially the FoxP2-expressing neurons in the KF and centro-lateral PB, have been shown to regulate ventilation^24^, and also drive cold-defense pathways that maintain core temperature^25,26^. These circuits, therefore, represent plausible sites where Oprm1–Adra2a co-activation could amplify respiratory and thermoregulatory depression during polysubstance exposure.

Despite this strong mechanistic rationale, no prior study has directly tested whether fentanyl and xylazine interact physiologically to produce additive or synergistic suppression of autonomic function. This gap is particularly consequential because xylazine is not an opioid and thus falls outside standard overdose research frameworks, toxicology testing, and clinical response protocols. Here, we address this gap by examining the physiological consequences of combined fentanyl and xylazine exposure in vivo in mice. We show that F+X produces markedly greater and more sustained reductions in respiratory rate and core body temperature than either drug alone, even at fentanyl doses typically considered safe. These findings provide the first direct experimental evidence that fentanyl and xylazine interact to depress vital autonomic functions, offering a mechanistic explanation for the disproportionate lethality observed in human overdose cases. More broadly, our results establish a framework for understanding how distinct inhibitory receptor systems may converge to amplify respiratory and thermoregulatory failure during emerging forms of polysubstance exposure.

## Material and Methods

### Animals

Adult male C57BL/6J mice (n = 12) were used. Mice were group-housed prior to surgical implantation and singly housed thereafter under controlled conditions (21–23°C; 40–60% humidity; 12 h light/dark cycle) with ad libitum access to food and water. Additonal, mice were surgically implanted with EEG/ EMG wire electrodes and telemetry devices (DSI, PhysioTel; n=3), as described in our published studies^29,33–35^. Meloxicam SR (4 mg/kg) was administered for postoperative analgesia. At study completion, mice were deeply anesthetized and perfused with 10% neutral buffered formalin; brains were collected for histology.

All procedures were approved by the Beth Israel Deaconess Medical Center IACUC (Protocol #032-2020-23) and complied with NIH guidelines.

### Drug Administration Protocol

Mice received intraperitoneal injections of saline or the following drugs (details in Key Resources table):

- Saline (S)-sterile 0.9% sodium chloride
- Fentanyl (F): 100, 350, and 600 μg/kg
- Xylazine (X): 6 mg/kg
- Fentanyl + Xylazine (F+X): F-350μg/kg + X-6mg/kg

Injections occurred at 11:00 AM (ZT4) following 2 h acclimation in the plethysmograph (9–11 AM). Treatments were separated by ≥7 days.

### Data acquisition

*EEG/EMG and respiration:* The recording sessions were conducted for 4-6 h to obtain a continuous baseline, or with either of the drug challenges (S, F, X, or F+X) for EEG/EMG and respiration, after placing the mice in a plethysmograph. The drugs were injected after the mice acclimatized to the recording chamber and the plethysmograph. To avoid confounding carryover or sensitization effects, drug challenges were administered in a fixed order with at least a week between the two, and with the highest fentanyl dose and the F+X combination always performed last. Across all conditions, each mouse received a maximum of four injections.

### Core Body Temperature Measurements

We conducted a recording of core body temperature by a) thermal imaging and b) abdominal core temperature, continuously for a period of 24hours, after saline injection or after the drug challenges. These longer recording sessions were separate from those of sleep and respiration, as plethysmograph recordings were limited to only 6-8h.

Thermal Imaging: Mice with shaved backs were continuously monitored using FLIR infrared cameras. Quantitative thermal images captured vasomotor and thermoregulatory responses to F+X as shown in **Fig. 2a**, consistent with prior work^35–38^. *Telemetry:* Intraperitoneal DSI PhysioTel transmitters^35–38^, recorded core temperature continuously for 24 hours following saline or drug injection. These longer recording sessions of core body temperature were conducted separately from plethysmograph recordings, which could only be done maximally for 6h.

*RNAscope FISH for Oprm1 and Adra2a expression in the PB: 20-micron sections were processed in RNAase-free conditions for labeling of Oprm1 or* Adra2a mRNA alongside the cell identity markers: FoxP2 mRNA; Calca mRNA (CGRP) by in situ hybridization (ISH). We used *RNAscope Multiplex Fluorescent Kit* V2 (Catalog # 323100, Advanced Cell Diagnostics, Hayward, CA). The sections will be treated with hydrogen peroxide, then with the target retrieval reagent for 5 minutes. These will then be dehydrated, airdried, treated with protease, and incubated in RNAscope probes for Oprm1 (Mm-Oprm 1-C2, Catalog # 316921-C2) or Adra2a (Mm-Adra2a-C3; Catalog # 425341-C3) for 2 hours for probe hybridization. Sections will then be subjected to 3 amplification steps at 40°C followed by incubation in HRP-C1 and TSA plus Fluorescein (Catalog # NEL741001KT, Perkin Elmer, MA) for 30 minutes at 40°C to visualize (Channel 1 at 488nm) for FoxP2, incubation of sections in HRP-C2 and TSA plus Cy3, and then Cy5 to visualize OPRM1, and CGRP mRNA. A separate set of brain sections was incubated only in HRP-C3 in TSA plus Cy5 to visualize Adra2a mRNA. Final slides were dried and cover-slipped with an antifade mounting reagent.

### Quantification and Statistical Analysis

#### Respiratory Analysis

Respiratory signals were analyzed using Spike2 Resp80t scripts (CED, UK). Trials were selected by investigators blinded to treatment for five steady-breath epochs during nonrapid eye movement sleep (NREM) state. Breath-by-breath analysis generated:

- Respiratory rate (RR)
- Tidal volume (VT)
- Minute ventilation (MV)

Values were quantified across time following saline, fentanyl, xylazine, or F+X injection (see **Fig. 1b**). Comparisons of RR were made as a percentage of their pre-injection values across time and treatment, and statistically compared, and plotted using SigmaPlot 14.5.

**Figure 1.**
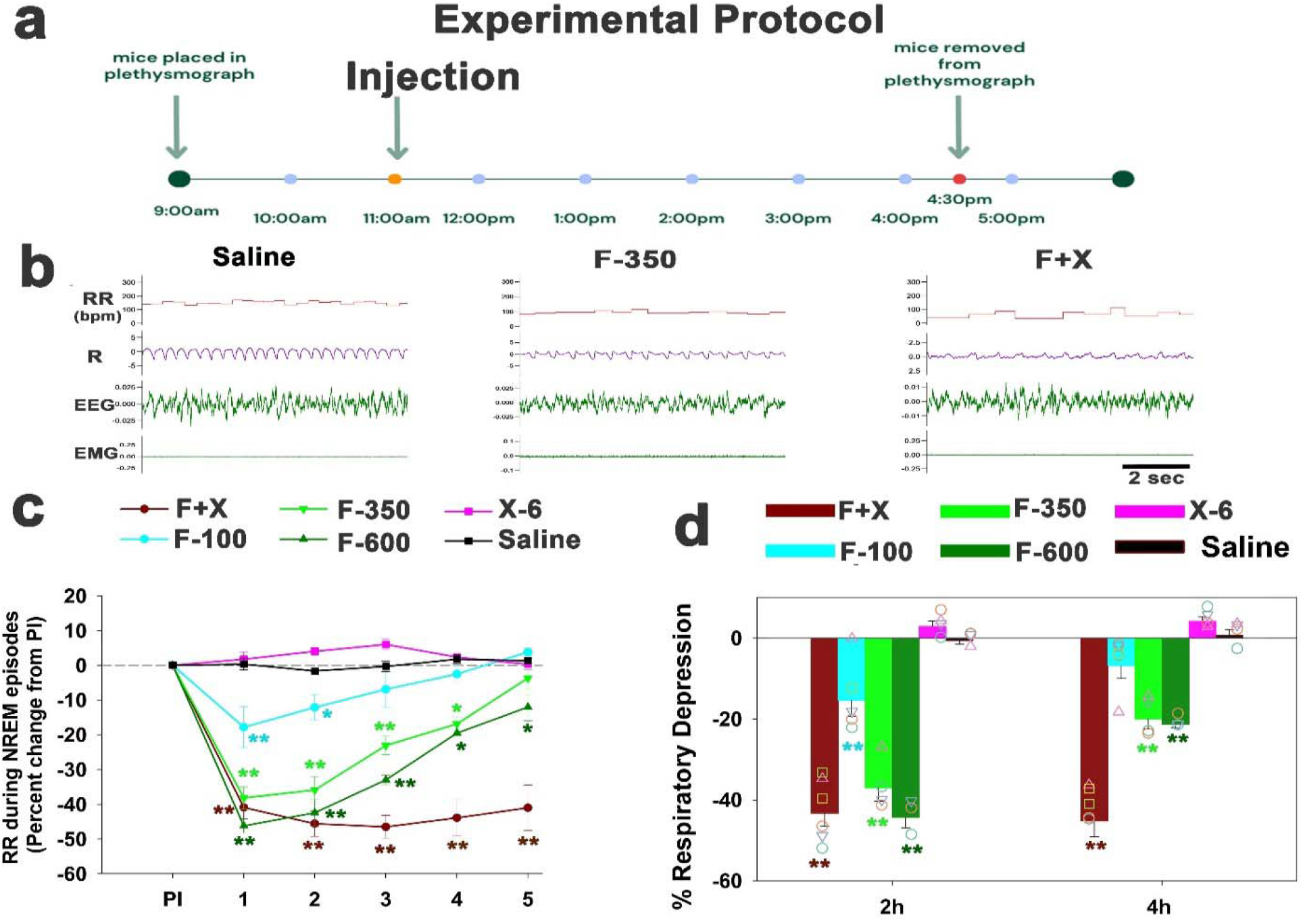
**a. Experimental protocol**. To test whether fentanyl (F) and xylazine (X) interact physiologically to suppress autonomic function, we quantified respiratory and thermoregulatory responses in freely behaving C57BL/6J mice (n = 12). Mice were implanted for EEG/EMG recording and monitored in whole-body plethysmography chambers. At ZT4, animals received intraperitoneal injections of fentanyl (100–600 μg/kg), xylazine (6 mg/kg), or the combination (F+X). Respiratory rate (RR), EEG/EMG, and core temperature were continuously recorded. **b. Representative physiological traces**: Example plethysmography, EEG, and EMG recordings illustrating the effects of saline, fentanyl (350 μg/kg; F-350), and F+X on respiratory pattern and behavioral state. F+X produces a deeper and more sustained suppression of respiratory activity compared to fentanyl (F-350) alone. **c. F+X produces prolonged opioid**-**induced respiratory depression (OIRD):** Fentanyl alone (100–600 μg/kg) induced a dose-dependent reduction in RR lasting 2–4 h, whereas xylazine alone (6 mg/kg) produced no significant change. In contrast, F350 μg/kg + X6 mg/kg (F+X) caused >40% depression of RR that remained significantly reduced for 5 h. **d. Quantification of respiratory depression:** Percent RR depression was calculated relative to each animal’s pre-injection baseline and assessed at 2 h and 4 h post-injection. Individual mouse values are shown as symbols overlaid on bar graphs. F+X produced significantly greater and more persistent RR suppression than either drug alone at both time points. Statistical comparisons were performed using two-way ANOVA (factors: treatment × time; n=4-6/ group), with post-hoc comparisons to saline. *P < 0.05; **P < 0.001.

**Figure 2.**
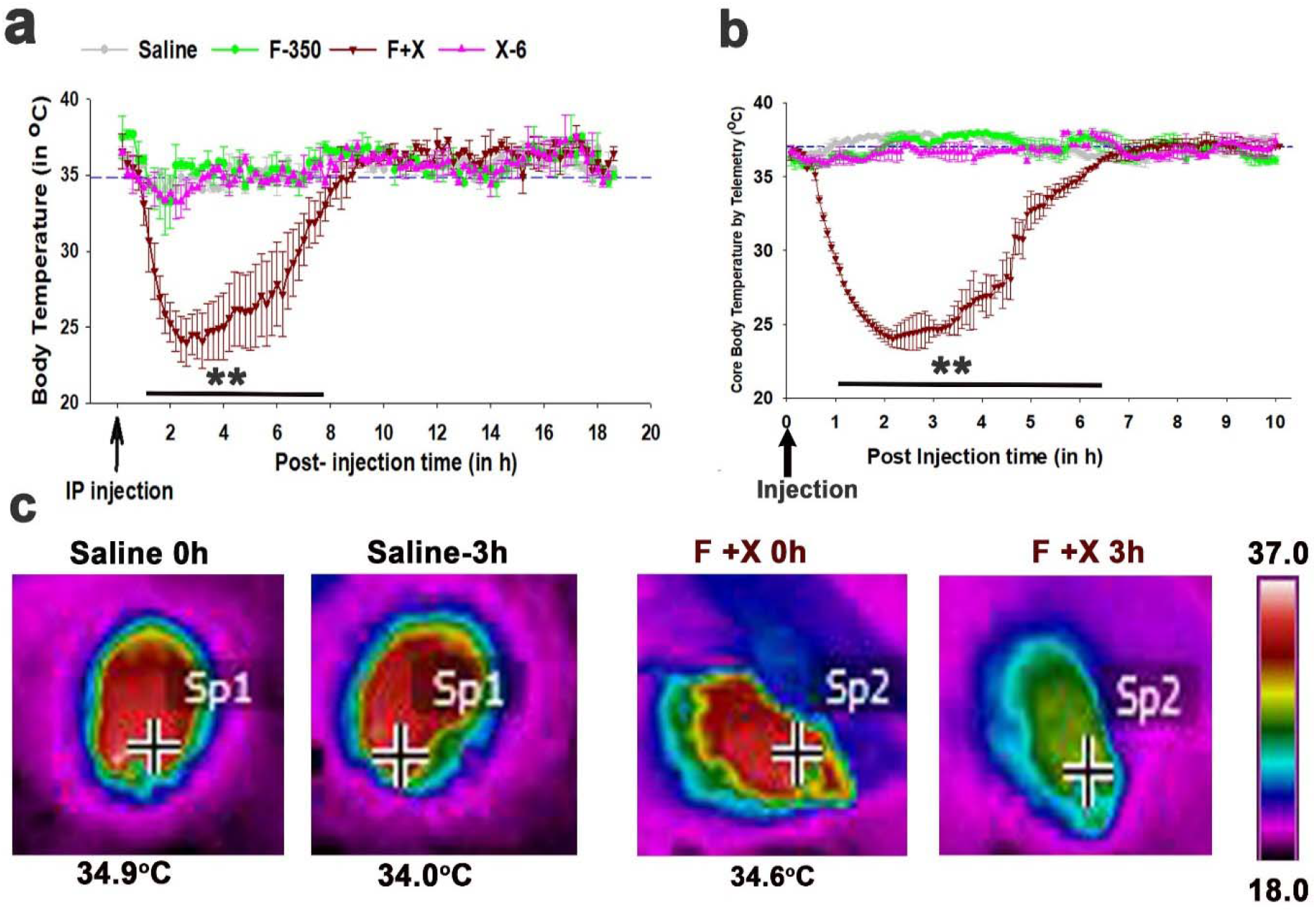
**a. Effect on body temperature:** Fentanyl + xylazine (F350 μg/kg + X6 mg/kg; F+X) induced a marked hypothermia, reducing both the temperature measured at the surface of the skin and core body temperature by 9-10°C. The hypothermic response induced by F+X lasted for ~7 h post-injection. In contrast, fentanyl (F-350) and X-6 alone had no effect on body temperature. **b. Core body temperature data collected using telemetry**, plotted similarly to that in a, also confirmed a reduction in body temperature by F+X, lasting for nearly 7-8 h, while F or X alone did not affect the body temperature. Data points recorded every 5 sec and averaged over 5 minutes are plotted in the graph for n=3 for each treatment. **c. Infrared thermal imaging:** Representative FLIR images from saline- and F+X-treated mice. At 3 h post-injection, F+X produced the maximal body temperature drop (~10 °C). Cross-marks indicate the precise points used for temperature recording. Statistical analysis: Temperature changes were compared to each animal’s pre-injection (PI) baseline using two-way ANOVA (factors: treatment × time; n = 4–6/ group). *P < 0.05; **P < 0.001 versus PI.

Temperature Analysis: Thermal imaging data were extracted from FLIR software every 10 minutes, by placing the cross hairs between the shoulder blades just below their neck. Telemetry data were analyzed using DSI acquisition software. Temperature traces were aligned to injection time and averaged across animals for every 5 minutes. Averaged core body temperature was statistically compared across time and the treatment groups, and plotted using SigmaPlot 14.5.

### Statistics

All statistical analyses were performed using SigmaPlot 14.5. Data are presented as mean ± SEM unless otherwise noted. For comparisons across treatment conditions (S, F, X, F+X) were performed using one or two-way ANOVA, followed by Holm–Sidak post hoc tests for pairwise comparisons. When time-course data were analyzed (e.g., RR, VT, MV, or temperature over time), two-way ANOVA (treatment × time) was used. Statistical significance was defined as p < 0.05.

### Data and Code Availability

- Respiratory analysis scripts (Spike2 Resp80t) are commercially available from CED.
- All datasets generated during this study are available upon reasonable request.

## KEY RESOURCES TABLE

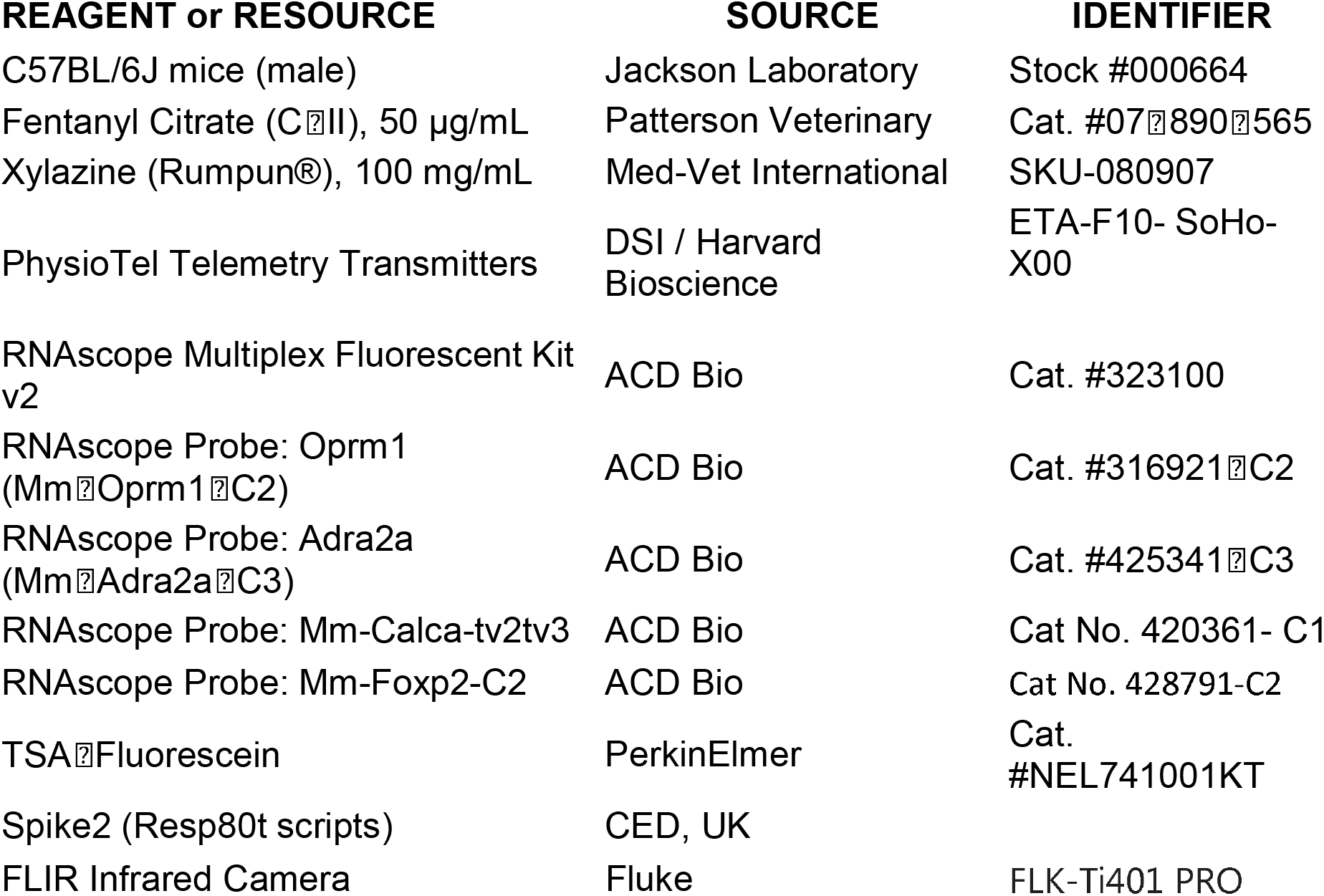

## Results

### Combined fentanyl and xylazine administration produces sustained respiratory depression

To test whether fentanyl (F) and xylazine (X) interact physiologically to suppress autonomic function, we quantified respiratory and thermoregulatory responses to each drug alone and in combination in freely behaving C57BL/6J mice (n = 12). Respiration was recorded using whole-body plethysmography following intraperitoneal (i.p.) injection of 0.9% saline, fentanyl (100–600 μg/kg), xylazine (6 mg/kg; X-6), or fentanyl (350 μg/kg) with xylazine (6 mg/kg) (F+X) at ZT4 (**Fig. 1a**). We selected a xylazine dose of 6 mg/kg because prior studies demonstrate that xylazine range (3–10 mg/kg) lies at the upper end of the behavioral range reported by Acosta-Mares et al.^2,39^ and well below the high-dose lethality range (≥30 mg/kg)^9^, while avoiding confounding sedative from xylazine alone. EEG/EMG was also recorded using implanted electrodes as sleep/wake state is necessary to guide respiration analysis. Accordingly, respiratory rate (RR) was analyzed during NREM sleep, as it provides the most stable and consistent signal, across the 5 h post-injection period (**Fig. 1b**).

Fentanyl alone produced a robust, dose-dependent respiratory depression lasting 2–5 h (F_5,120_ = 9.4; P < 0.001) compared to saline (n=4) (**Fig. 1c**). Low-dose fentanyl (F-100; n= 5) caused a modest RR reduction (15.4 ± 3.9%) limited to 2 h (F_15, 132_ = 15.6; P < 0.001), whereas higher doses (F-350, n= 4; and F-600, n=3) induced marked suppression at 1h (37% and 44%, respectively; F_5,120_ = 7.9 and 8.9; P < 0.001) that partially recovered by 5 h (**Fig. 1d**). Xylazine alone (n= 4) did not alter RR at any time point. In contrast, co-administration of fentanyl (350 μg/kg) with xylazine (6 mg/kg) (F+X, n=6) produced a prolonged respiratory depression persisting throughout the 5 h recording period (F_5,120_ = 142.5; P < 0.001). RR suppression for the first two hours (2h, 43.3%) was comparable to F-600 alone but remained >40% at 4–5 h, indicating sustained inhibition beyond the pharmacodynamic window of either drug alone (**Fig. 1d**). As xylazine alone has no effect on respiratory rate, the interaction between fentanyl and xylazine reflects pharmacodynamics potentiation likely through the interactions between Oprm1 and Adra2a receptor pathways rather than additive effects of two independently depressant agents.

### Fentanyl–xylazine co-exposure induces prolonged hypothermia

To determine whether F+X also affects thermoregulation, we measured core body temperature using infrared thermography (FLIR) and implanted telemetry probes (DSI PhysioTel). FLIR analysis revealed significant hypothermia in F+X-treated mice (n=4) (F_5,354_ = 82.03; P < 0.001) lasting up to 7 h post-injection (**Fig. 2a**). Neither fentanyl (n=3) nor xylazine (n=3) alone altered core body temperature. This was also confirmed by recording core body temperature with implanted DSI devices and telemetry in additional set of mice (n=3), in which F+X-induced hypothermia lasted for 7h (F_3, 1387_ = 115.2; P < 0.001) (**Fig. 2b**). Representative FLIR images (**Fig. 2c**) illustrate the marked drop in the body temperature at 3 h post-injection, notably following the nadir of RR suppression at 1-2 h post injection. Telemetry recordings confirmed these findings, showing sustained reductions in core temperature following F+X but not saline or fentanyl conditions. The time course of hypothermia and respiratory depression suggests coordinated suppression of autonomic circuits controlling ventilation and thermogenesis.

### Fentanyl–xylazine lethality and receptor co-expression in brainstem circuits

The F+X combination reduced survival to 58.8% (7/12) within 5 h post-injection, whereas xylazine alone and lower fentanyl doses (100–350 μg/kg) did not a cause mortality; high-dose fentanyl (600 μg/kg) was lethal in 2/12 animals (**Fig. 3a**). Animals were monitored throughout and after drug administration for signs of severe autonomic compromise. One mouse in the F+X group died 4h post-injection; however, the physiological data acquired prior to were technically robust and were therefore retained for analysis and included in **Fig. 1c–d**. Excluding this animal would have biased the dataset toward survivors and underestimated the severity of F+X toxicity; therefore, its pre-mortem data were retained. Since combined fentanyl–xylazine exposure produced profound but often reversible hypothermia, transient reductions in core temperature below 24 °C were not by themselves reason enough for euthanasia. However, seven of twelve F+X-treated mice failed to recover, exhibiting a progressive decline in core temperature to <24 °C during the 3-4h post-injection window, followed by a prolonged state of severely diminished, labored respiration that culminated in death. The selective lethality of the F+X combination underscores a potentiating autonomic depressant effect that exceeds the toxicity profiles of fentanyl or xylazine individually.

**Figure 3.**
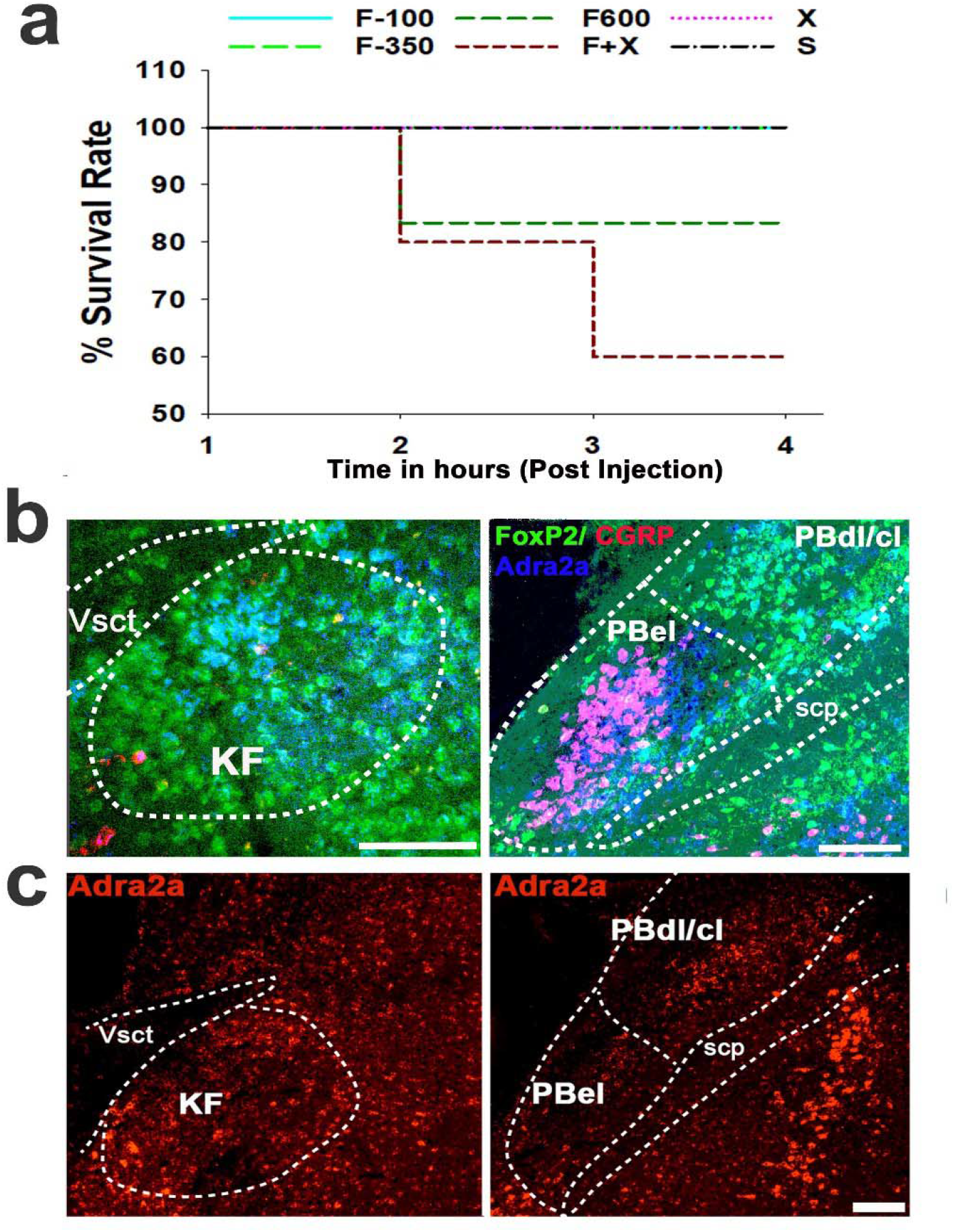
**a. Survival analysis across treatments:** Kaplan–Meier survival curves showing differential lethality across drug conditions. The fentanyl–xylazine combination (F350 μg/kg + X6 mg/kg; F+X) reduced survival to 58.8% (7/12) within 5 h post-injection. Xylazine alone and lower fentanyl doses (100–350 μg/kg) did not affect survival, whereas high-dose fentanyl (600 μg/kg) resulted in mortality in 2/12 animals. **b. Validation of Oprm1 expression in PB/KF neurons**: RNAscope in situ hybridization confirming Oprm1 mRNA expression in FoxP2-positive and CGRP-positive neurons within the parabrachial (PB) and Kölliker-Fuśe (KF) complex. Scale bar: 100 μm. **c. Distribution of Adra2a receptors in PB and KF:** RNAscope labeling of Adra2a mRNA showing receptor-expressing neurons across PB subnuclei, including the external lateral (el) and center-lateral (cl) regions, as well as the medial PB (mPB). The KF area also showed the presence of Adra2a. Anatomical landmarks include the ventral spinocerebellar tract (Vsct) and superior cerebellar peduncle (scp). Scale bar: 100 μm.

To identify potential neural substrates for interaction between opioid and α_2_-adrenergic receptors, we examined Oprm1 and Adra2a expression within the PB and KF nuclei that regulate ventilation^23,24^. Using RNAscope on mouse brainstem sections, we confirmed and extended prior single-cell and spatial transcriptomic findings^22^ by demonstrating that Oprm1 is expressed in both FoxP2- and CGRP-positive PB/KF neurons (**Fig. 3b**), and that Adra2a is present within PB/KF FoxP2 populations which have been shown to regulate ventilation^24^ by our group (**Fig. 3c**). These findings establish that both receptor systems are poised to influence PB/KF output, but they do not resolve whether Oprm1 and Adra2a are co-expressed within the same neurons or segregated across parallel FoxP2 subpopulations. Accordingly, the proposed dua-lreceptor convergence model should be interpreted as a mechanistic hypothesis grounded in anatomical overlap, rather than evidence of demonstrated cellular co-localization.

## Discussion

Our findings demonstrate that fentanyl and xylazine interact physiologically to produce sustained suppression of respiratory and thermoregulatory function, consistent with convergent inhibition of brainstem^32,40–42^ and hypothalamic^26,37,43^ autonomic circuits.

Whereas fentanyl alone produced dose-dependent respiratory depression with partial recovery over several hours and did not alter body temperature, co-administration with xylazine markedly prolonged the respiratory deficit and induced profound hypothermia. The absence of respiratory or temperature effects from xylazine alone indicates that the interaction is not simply additive but reflects pharmacodynamic convergence between μ-opioid and α_2_-adrenergic receptor systems^12,13^, each of which independently regulates arousal, respiratory rhythmogenesis, and thermogenic drive. Only a few studies^8,39^ have previously tested the potentiation of fentanyl-induced brain hypoxia via xylazine, that was partially reversed by naloxone. However, this result provides direct experimental evidence that simultaneous engagement of these inhibitory GPCR pathways produces a level of autonomic suppression that exceeds the effects of either drug alone.

Although F+X co-exposure produced profound hypothermia, several observations indicate that cooling is not initiating respiratory depression^44^. Respiratory rate reached its nadir within the first hour, whereas core temperature continued to decline for several hours thereafter, demonstrating a temporal dissociation in which hypothermia developed downstream of the primary depressant mechanism. Moreover, fentanyl alone produced substantial respiratory depression without altering body temperature, indicating that μ-opioid–mediated inhibition of respiratory circuits is sufficient to suppress breathing in the absence of hypothermia. Finally, the magnitude of respiratory depression in the F+X condition exceeded what would be expected from metabolic suppression due to the body cooling ^45^, as mild–moderate hypothermia primarily reduces ventilatory demand through decreased CO_2_ production rather than directly inhibiting respiratory rhythm generation^46–48^.

Since xylazine alone did not alter respiratory rate or core body temperature, the interaction observed in the F+X condition reflects pharmacodynamic potentiation rather than classical additivity or synergy. Despite the use of the term ‘synergistic’ to describe the opioid and α_2_-adrenergic receptor interactions in earlier reports^9,13–15^ we do not apply it here because synergy must be demonstrated through formal interaction analyses-such as isobolographic or dose equivalence testing^9^—which were not part of the present experimental design. Instead, our data support an additive or potentiating interaction, in which simultaneous activation of μ-opioid and α_2_-adrenergic receptor pathways produce a level of autonomic suppression greater than either drug alone.

Together, these findings support a model in which hypothermia acts as a secondary amplifier of respiratory depression induced by F+X rather than as the primary cause of the breathing deficit.

The prolonged depression of respiratory and thermoregulatory function observed during F+X exposure is consistent with convergent inhibition within PB/KF, which integrate chemosensory, arousal, and thermogenic signals. μ-opioid (Oprm1) -expressing PB/KF neurons are principal mediators of opioid-induced respiratory depression^32,40–42^. These nuclei integrate respiratory^24^, arousal^29,49^, and thermoregulatory^26^ information and project to both the pre-Bötzinger complex preBötC^24^, which generates inspiratory rhythm, and hypothalamic centers such as the dorsomedial hypothalamus (DMH)^26^ and the preoptic area (POA), which regulate thermogenesis.

Although our RNAscope data demonstrate that Oprm1 and Adra2a transcripts both are present within PB/KF FoxP2 populations, but these experiments cannot determine whether the receptors are co-expressed within the same neurons or distributed across parallel subpopulations that converge onto shared downstream targets. Accordingly, the dua-lreceptor convergence model we propose here should be viewed as a mechanistic framework supported by anatomical overlap and the robust physiological interaction, rather than a claim of demonstrated cellular co localization. Future studies using ce-lltype-specific manipulations will be required to resolve whether fentanyl and xylazine act on single dua-lexpressing neurons or through coordinated inhibition across distinct PB/KF microcircuits. Importantly, this limitation does not alter the central conclusion that simultaneous engagement of Oprm1 and Adra2a pathways produces additive and sustained autonomic depression in vivo.

These findings have direct implications for understanding the disproportionate lethality of fentanyl–xylazine mixtures in humans. Xylazine is not an opioid and therefore falls outside standard overdose response protocols, toxicology testing, and naloxone-based reversal strategies. The prolonged respiratory depression and profound hypothermia observed in our study mirror clinical reports of deep unresponsiveness and naloxone-insensitive hypoventilation in individuals exposed to F+X^3,9^. By demonstrating that μ-opioid and α_2_-adrenergic receptor pathways interact to produce sustained autonomic suppression, our results provide a potential mechanistic basis for why standard opioid antagonism may be insufficient in F+X overdose, and highlight the need for adjunctive therapeutic strategies targeting α_2_-adrenergic signaling or downstream autonomic circuits.

### Limitations and Future Directions

First, our experiments focused on acute responses; chronic or repeated fentanyl–xylazine exposure may engage adaptive mechanisms that may alter receptor sensitivity or circuit connectivity, which were not studied in the present study. Second, our RNAscope data cannot determine whether both Oprm1 and Adra2a receptors reside within the same neurons or in parallel FoxP2 subpopulations that converge onto shared downstream targets. However, importantly, the physiological interaction we observe— with markedly prolonged respiratory and thermoregulatory suppression during fentanyl–xylazine co-exposure—requires only that both receptor pathways inhibit PB/KF output, not that they necessarily be co-localized within single neurons. Future ce-lltype-specific manipulations will be essential to resolve whether the interaction arises from true co-expression or coordinated inhibition across distinct PB/KF microcircuits. Future studies using ce-lltype-specific manipulations and in vivo circuit mapping will be conducted to determine whether Oprm1- and Adra2a-expressing FoxP2 neurons act cooperatively or through parallel pathways. Finally, translational studies are needed to test whether α_2_-adrenergic antagonists or targeted activation of PB/KF output can mitigate fentanyl–xylazine–induced autonomic failure. Addressing these questions will refine our understanding of polysubstance-induced respiratory depression and inform therapeutic strategies for emerging synthetic opioid mixtures.

## Acknowledgments

We thank Quan Ha and Sam Sailesh for their excellent technical support and assistance with data scoring.

## Authorship Contributions

NL-Data collection and manuscript writing; JDL-data analysis and brain tissue processing; SSB-mouse breeding program and RNA scope processing; NM-Data acquisition, analysis, and manuscript writing; SK-experimental design and conceptualization, data collection, analysis, and manuscript writing.

